# Endothelium-Dependent Vasodilation is Impaired in Chronic Spinal Cord Injury and is Associated with Oxidative Stress

**DOI:** 10.64898/2026.05.27.728335

**Authors:** Andrew J. Park, Christopher A. DeSouza, Genevieve Madera, Clare Morey, Megan Summers, Vinicius P Garcia, Auburn R. Berry, Samuel T. Ruzenne, Noah M. DeSouza, Joshua P. Holzer, Alexa Deitemeyer, Jared J. Greiner, Brian L. Stauffer

## Abstract

**Background:** Individuals with spinal cord injury (SCI) experience accelerated atherosclerotic cardiovascular disease that is not fully explained by traditional risk factors. Endothelial dysfunction is a key mechanism in atherosclerosis. We tested the hypothesis that endothelium-dependent vasodilation is impaired in adults with SCI and is due, at least in part, to oxidative stress.

**Methods:** Twenty-four adults (age:19-58 yr) free of overt cardiometabolic disease were studied: 12 non-injured adults (9 M/3 F) and 12 adults with chronic SCI (8 M/4 F; time since injury 1.5 – 25 years). Forearm blood flow was determined (FBF; via strain-gauge plethysmography) in response to intra-arterial infusion of acetylcholine and isoproterenol in the absence and presence of the antioxidant vitamin C as well as the FBF response to sodium nitroprusside.

**Results:** Adults with SCI demonstrated significantly lower vasodilator response to acetylcholine (from 4.1±0.6 to 10.7±2.6 mL/100 mL tissue/min vs 4.1±1.1 to 15.7±3.4 mL/100 mL tissue/min) and isoproterenol (4.0±0.6 to 11.2±2.2 mL/100 mL tissue/min vs 4.3±1.0 to 15.0±2.6 mL/100 mL tissue/min) compared with non-injured adults. FBF response to sodium nitroprusside was not significantly different between the groups. Co-infusion of vitamin C significantly increased the vasodilator response to acetylcholine (~45%) and isoproterenol (~25%) in the adults with SCI to levels comparable with non-injured adults.

**Conclusions:** Chronic SCI is associated with endothelial-dependent vasodilator dysfunction. Impaired vasodilation across two distinct endothelial agonists suggests that chronic SCI is associated with endothelial dysfunction not confined to a specific receptor or intracellular signaling pathway.

Moreover, oxidative stress is a contributing factor underlying SCI-related endothelial vasodilator dysfunction.

NCT06443151

**CLINICAL PERSPECTIVE:**

- The novel finding of this study is that individuals with SCI demonstrate impaired endothelial vasodilator function in absence of traditional cardiovascular risk factors.
- Oxidative stress is a contributing factor to SCI-related endothelial vasodilator dysfunction.
- Future studies are needed to determine the efficacy of therapeutic interventions, either lifestyle or pharmacologic, in improving endothelial function in order to mitigate the elevated ASCVD risk after SCI.

## INTRODUCTION

Individuals living with spinal cord injury (SCI) experience a markedly accelerated burden of atherosclerotic cardiovascular disease (ASCVD), with myocardial infarction and ischemic stroke occurring more frequently and earlier in life than in adults without SCI. Several independent epidemiological studies demonstrate an overall higher incidence of major adverse cardiovascular events in individuals with SCI compared with age-, sex-, and comorbidity-matched controls.^1–3^ Alarmingly, the study by Yoo et al 2024, not only demonstrated a 2-fold increased incidence of myocardial infarction and heart failure, but much earlier age of onset ^3^. Additionally, increased atherosclerotic burden has been detected among individuals with SCI, otherwise classified as low risk by traditional cardiovascular risk factors.^4–8^ Multiple assessment methods, including carotid intima-media thickness measurement, nuclear imaging studies, and coronary artery calcium scoring via computed tomography, consistently demonstrate elevated atherosclerotic burden in this population. These multimodal approaches provide converging evidence that individuals with SCI face heightened risk, even when conventional risk factors might suggest otherwise.

Endothelial dysfunction, particularly impaired endothelium-dependent vasodilation, is a central etiological component in the development of ASCVD, ^9–14^ occurring before histologic and/or angiographic evidence of atherosclerosis.^9,10,14–20^ While many of the cardiovascular consequences associated with SCI, such as myocardial infarction, heart failure, and stroke, have been attributed, at least in part, to endothelial vasomotor dysfunction ^21–23^, prior studies have been equivocal and difficult to interpret because of reliance on largely noninvasive approaches such as nonspecific circulating biomarkers, variable experimental conditions, and heterogeneous SCI cohorts.^2,24–29^ The latter limitation is an important consideration as the level and completeness of injury can differentially affect vascular function and ASCVD risk.^2,28^ Establishing whether endothelium-dependent vasodilation is impaired in adults with SCI and, if so, identifying potential mechanisms is vital for developing effective prevention and treatment strategies for ASCVD in this high-risk population.

Accordingly, the experimental aim of the present study was to determine if endothelium-dependent vasodilation is impaired in adults with SCI. We hypothesized that endothelium-dependent vasodilation is impaired in adults with SCI compared with non-injured adults. Moreover, the SCI-related reduction in endothelial vasodilator function is not limited to a specific endothelial receptor or signal transduction pathway and is due, at least in part, to increased oxidative stress. To test our hypotheses, we determined forearm vascular responses to various endothelial agonists that stimulate nitric oxide (NO)-mediated endothelium-dependent vasodilation via different cell surface receptors and intracellular signaling pathways in the absence and presence of the potent antioxidant, vitamin C.

## METHODS

### Subjects

Twenty-four young and midlife adults (age:19-58 yr) free of overt cardiovascular and metabolic disease were studied: 12 non-injured adults (9 M/3 F) and 12 adults with chronic cervical and high thoracic injuries (8 M/4 F; time since injury 1.5 – 25.13 years). As neurological level of injury may influence vascular function and outcome after SCI ^28^, only individuals with cervical and high thoracic injury determined by the International Standards for Neurological Classification of Spinal Cord Injury (ISNCSCI) were included in the study.^30^ Neurological level of injury (NLI) in the SCI group included T2: n=1; T3: n=1; T4: n=4; T6: n=2; T9 n=1; T10 n=1; T11 n=1; T12 n=1. Completeness of injury was defined by the American Spinal Injury Association (ASIA) Impairment Scale (AIS). AIS in the SCI group included: A: n=10; C: n=2.

All non-injured and SCI adults were free of overt cardiometabolic disease by medical history, physical examination, and hematologic evaluation. Fasting plasma lipid, lipoprotein, and glucose concentrations were determined using standard techniques. Body mass index was calculated as kg/m^2^ with body mass measured to the nearest 0.1 kg and height to nearest 0.1 meters.

### Metabolic Measurements

Fasting plasma lipid and lipoprotein, glucose and insulin concentrations were determined using standard techniques by the clinical laboratories affiliated with Craig Hospital and the Clinical Translational Research Center at the University of Colorado Boulder.

### Intra-arterial Infusion Protocol

All studies were performed between 7:00 am and 10:00 am after a 12-hour overnight fast in a temperature-controlled room. Under strict aseptic conditions a 5-cm, 20-gauge catheter was inserted into the brachial artery of the nondominant arm under local anesthesia (1% lidocaine). Heart rate and arterial pressure were continuously measured throughout the infusion protocol. Forearm blood flow (FBF) was measured in both the experimental (nondominant) and contralateral (dominant) forearm using strain-gauge venous occlusion plethysmography (D.E. Hokanson) as previously described by our laboratory.^31^ All FBF values are presented in milliliters per 100 milliliters of forearm volume per minute. Forearm volume was determined using the water displacement method.^32^ Following the measurement of resting blood flow for 5 minutes, endothelium-dependent vasodilatation was assessed by the FBF responses to incremental doses of acetylcholine (Bausch Health, Bridgewater, NJ) and isoproterenol (Isuprel, Abbott Laboratories, Abbott Park, IL). FBF response to sodium nitroprusside (Hospira, Lake Forest IL) was used to assess endothelium-independent vasodilatation. Acetylcholine was infused at rates of 4.0, 8.0, 16.0 µg/100 mL of forearm tissue/min, isoproterenol at 5.0, 10.0 and 20.0 ng/100 mL tissue/min, and sodium nitroprusside at 1.0, 2.0, and 4.0 µg/100 mL of forearm tissue/min. Each dose was infused for 5 minutes and sufficient time (~20 minutes) was provided to allow FBF to return to resting levels between drug infusions. To avoid an order effect the sequence of drug administration was randomized. Vitamin C was infused at a constant rate (12 mg/100 mL tissue/min) for 5 minutes. This vitamin C concentration has been shown to both protect human plasma from free radical-mediated lipid peroxidation ^33^ and improve endothelium-dependent vasodilation in conditions associated with oxidative stress.^34^ Vitamin C infusion was maintained at the same rate whilst the acetylcholine and isoproterenol dose-response curves were repeated in the same order as performed earlier.

### Statistical Analysis

Differences in subject characteristics and area under the curve data were determined by between groups analysis of variance (ANOVA). Differences in the FBF responses to each vasoactive drug in the absence and presence of vitamin C were determined by repeated measures ANOVA. When appropriate, the Sequential Holm-Bonferroni *post-hoc* method was performed to determine within-group differences at each concentration of vasoactive drug. There were no significant main effects of sex or sex interaction in the FBF responses to vasoactive agents, therefore the data were pooled and presented together. Total blood flow released across the forearm in response to each vasoactive agent was calculated as the incremental area under each curve using a trapezoidal model. All values are presented as mean±SD. Statistical significance was set *a priori* at P < 0.05.

## RESULTS

Selected subject characteristics are presented in Table 1. There were no significant differences in age, body mass, blood pressure, plasma lipid and lipoproteins or glucose and insulin concentrations between the groups. Resting forearm blood flow in the non-infused arm and mean arterial pressure remained constant throughout the infusion protocols and were not significantly different between the groups (data not shown).

**Table 1.** Selected Participant Characteristics.

| Variable | Non-injured<br>(n=12) | SCI<br>(n=12) |
| --- | --- | --- |
| Age, yrs | 32±12 | 33±10 |
| Sex, M/F | 9/3 | 8/4 |
| Time since injury, months | -- | 118.8±103.2 |
| Body mass, kg | 77.4±14.7 | 72.7±11.9 |
| BMI, kg/m<sup>2</sup> | 24.3±3.5 | 23.3±4.2 |
| Systolic BP, mmHg | 117±12 | 118±11 |
| Diastolic BP, mmHg | 69±10 | 71±11 |
| Total cholesterol, mg/dL | 153±20 | 164±31 |
| LDL-cholesterol, mg/dL | 85±22 | 93±28 |
| HDL-cholesterol, mg/dL | 48±16 | 50±13 |
| Triglycerides, mg/dL | 99±47 | 106±50 |
| Glucose, mg/dL | 89±10 | 83±10 |
| Insulin, µU/mL | 5.0±1.9 | 7.7±4.7 |

BMI, body mass index; BP, blood pressure; LDL, low-density lipoprotein; HDL, high-density lipoprotein. Values are mean±SD.

Figure 1 shows the FBF responses to acetylcholine, isoproterenol, and sodium nitroprusside in the non-injured and SCI groups. The adults with SCI demonstrated a markedly blunted vasodilator response to both acetylcholine and isoproterenol. The increase in FBF to acetylcholine was ~30% less (P<0.01) in the SCI (from 4.1±0.6 to 10.7±2.6 mL/100 mL tissue/min) compared with the non-injured (from 4.1±1.1 to 15.7±3.4 mL/100 mL tissue/min) group. As a result, total FBF to acetylcholine (area under the FBF curve) was ~45% lower in the adults with SCI (45±16 mL/100 mL tissue) than the non-injured (83±23 mL/100 mL tissue) adults. In response to isoproterenol, FBF was ~25% lower (P<0.01) in the SCI (4.0±0.6 to 11.2±2.2 mL/100 mL tissue/min) versus non-injured (4.3±1.0 to 15.0±2.6 mL/100 mL tissue/min) group. Consequently, total FBF to isoproterenol was lower (~35%: P<0.01) in the SCI (47±2 mL/100 mL tissue) compared with non-injured (71±25 mL/100 mL tissue) adults. There were no significant correlations between time since injury and total FBF responses to either acetylcholine (r=-0.37; P=0.25) or isoproterenol (r=-0.36; P=0.24) in the SCI group. FBF responses to sodium nitroprusside were not significantly different (P=0.52) between the non-injured (from 4.5±1.2 to 16.6±2.6 mL/100 mL tissue/min) and SCI (from 4.1±1.0 to 15.4±3.4 mL/100 mL tissue/min).

**Figure 1.**
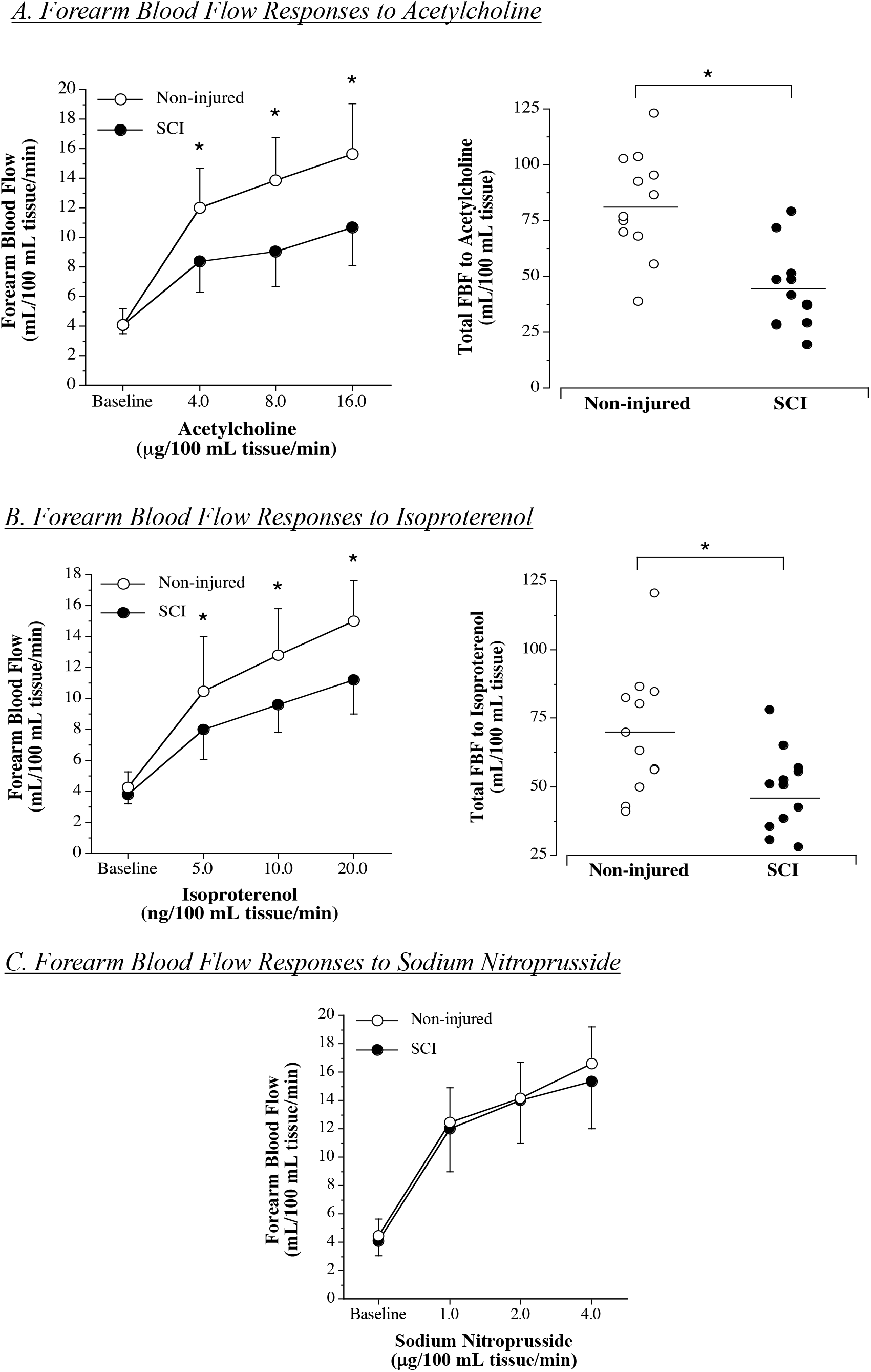
Forearm blood flow responses and total forearm blood flow (area under the curve) to acetylcholine (A), isoproterenol (B) and sodium nitroprusside (C) in the non-injured adults and adults with SCI. Values are mean ± SD. For area under the curve the mean value is denoted. *P<0.05 vs non-injured.

Figure 2 shows the vasodilatory response to the co-infusion of vitamin C with acetylcholine and isoproterenol in the adults with SCI. Vitamin C significantly increased (P<0.01) the FBF response (from 4.1±0.6 to 15.1±2.1 mL/100 mL tissue/min) and total FBF (83±17 mL/100 mL tissue) to acetylcholine (~45% and ~85%, respectively) in the adults with SCI. Similarly, the co-infusion of vitamin C significantly increased (P<0.01) the FBF response (from 4.1±0.7 to 13.9±3.1 mL/100 mL tissue/min) and total FBF (65±16 mL/100 mL tissue) to isoproterenol (~25% and ~40%, respectively) in the adults with SCI. The vasodilatory responses to both acetylcholine + vitamin C and isoproterenol + vitamin C in the adults with SCI were not significantly different from the vasodilatory response to acetylcholine and isoproterenol in the non-injured group.

**Figure 2.**
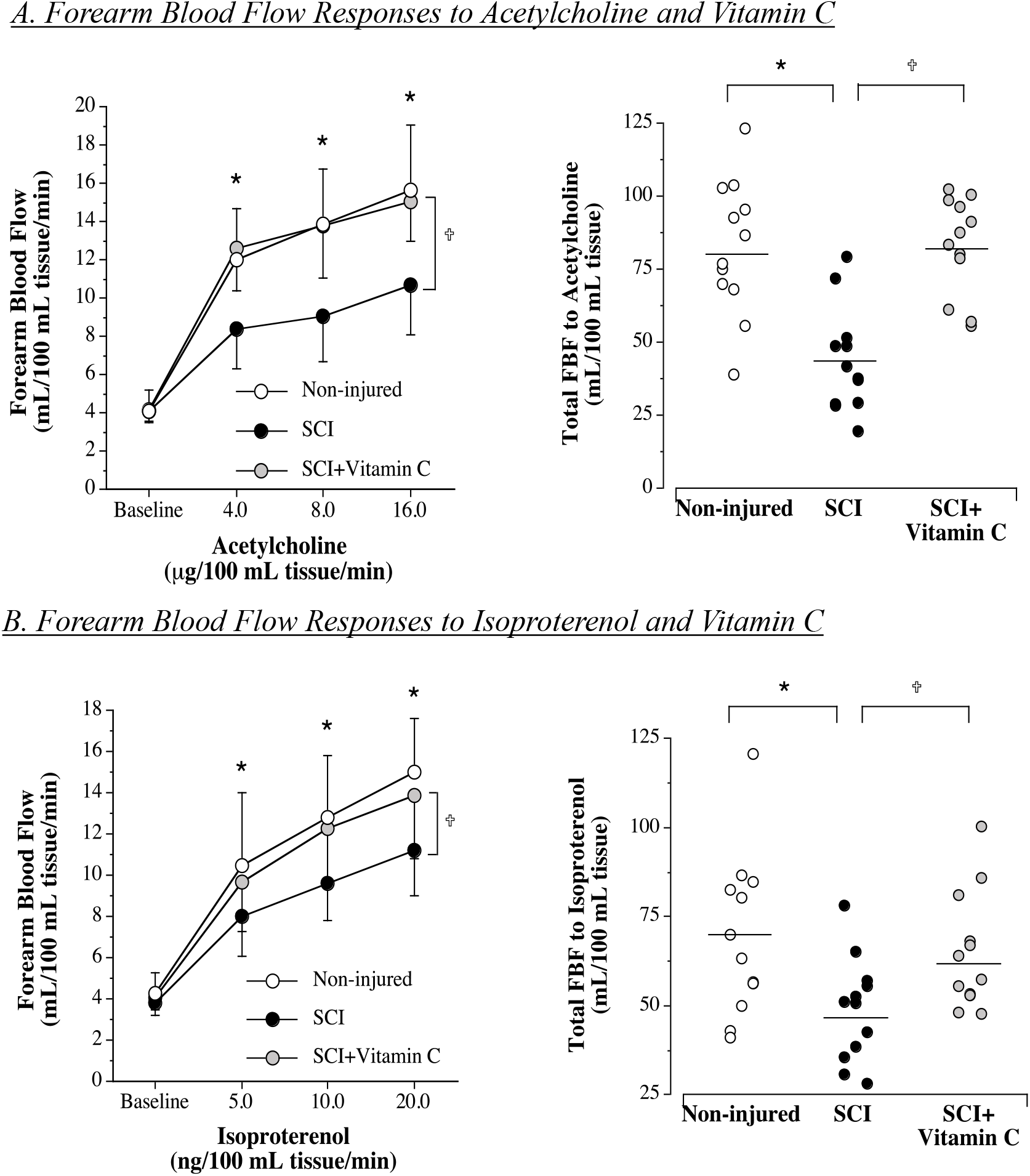
Forearm blood flow responses and total forearm blood flow (area under the curve) to acetylcholine (A) and isoproterenol (B) in the absence and presence of the antioxidant vitamin C in the adults with SCI. Values are mean ± SD. For area under the curve, the mean value is denoted. *P<0.05 vs non-injured; †P<0.05 vs saline within SCI adults.

## DISCUSSION

The novel and seminal finding of the present study is that chronic SCI is associated with profound endothelial-dependent vasodilator dysfunction. Indeed, we demonstrate for the first time that endothelium-dependent vasodilation in response to both acetylcholine and isoproterenol is significantly impaired, whereas endothelium-independent vasodilation to sodium nitroprusside is preserved in adults with SCI. Impaired vasodilation across two distinct endothelial agonists suggests that chronic SCI is associated with general dysfunction in agonist-mediated endothelium-dependent vasodilation that is not confined to a specific receptor or intracellular signaling pathway. Moreover, the results of the present study suggest that impaired endothelium-dependent vasodilation in adults with chronic SCI is due, at least in part, to oxidative stress.

To comprehensively assess the influence of SCI on endothelium-dependent vasodilation in conscious humans, we determined the FBF responses to two distinct endothelial agonists that activate endothelial nitric oxide synthase and, in turn, NO production; but act via different endothelial cell surface receptors, intracellular membrane-bound G proteins, and signal transduction pathways.^35^ Acetylcholine activates muscarinic receptors coupled to intracellular membrane-bound pertussis toxin-sensitive G protein, which stimulates the phospholipase C-phosphatidylinositol-Ca^2+^ signaling pathway. This leads to increased intracellular calcium and activation of eNOS.^35^ In contrast, isoproterenol activates β-adrenoceptors stimulating the adenylyl cyclase pathway resulting in increased cAMP and activation of eNOS.^36,37^ Herein we demonstrate that regardless of the agonist administered, endothelium-dependent vasodilation was significantly lower in adults with SCI compared with their non-injured peers. In addition, the magnitude of impairment in vasodilation was similar amongst the agonists (~25-30%) supporting the notion of endothelial vasomotor dysfunction with a common underlying cause. Indeed, although acetylcholine and isoproterenol activate endothelium-dependent vasodilation via distinct receptor-mediated signaling pathways they converge at endothelial nitric oxide synthase (eNOS)-dependent NO production. Thus, a reduction in eNOS activation resulting in less NO bioavailability would be a logical mechanism underlying the global impairment in endothelial vasodilator capacity in the adults with SCI. Supporting this postulate is our finding that the FBF response to sodium nitroprusside was not significantly different between the non-injured and SCI adults indicating that diminished NO bioavailability, rather than altered vascular smooth muscle sensitivity to NO, is a central factor underlying the observed vasodilator impairment after SCI.

Oxidative stress has deleterious effects on endothelial function contributing to vasomotor dysfunction.^38^ Data from our laboratory ^39^ and others ^38^ in non-injured adults have demonstrated that oxidative stress is associated with impaired endothelium-dependent vasodilation and is an important underlying factor in vascular dysfunction and disease.^40^ SCI is known to induce chronic oxidative stress ^41–43^. Polymorphonuclear neutrophil production of reactive oxygen species, particularly hydrogen peroxide, is markedly and persistently higher after SCI.^42,43^ In addition, antioxidant defense mechanisms such as cysteine and glutathione metabolism are diminished in adults with SCI.^41^ The consequences of increased oxidative stress after SCI are not completely understood. It has been suggested that heightened oxidative damage to cells of the vasculature and immune system may underlie several disease processes in adults with SCI.^44–46^ Intra-arterial administration of vitamin C at pharmacological doses, as used in the present study, is a well-validated technique to determine the impact of oxidative stress on endothelial vasodilator function in vivo. ^38,39^ Herein we demonstrate, for the first time, that co-infusion of vitamin C significantly enhances the endothelial vasodilator response to both acetylcholine and isoproterenol in adults with SCI to levels comparable with non-injured adults. As such, oxidative stress appears to be a major contributor to endothelial vasodilator dysfunction after SCI. Moreover, the fact that vitamin C improved the vasodilator response to both endothelial agonists provides additional support for impaired eNOS-dependent NO production as a central mechanism underlying the SCI-related dysfunction in endothelium-dependent vasodilation. It is well established that oxidative stress can severely compromise eNOS activity, and in turn, NO production through a variety of mechanism such as: oxidizing and neutralizing tetrahydrobiopterin a crucial cofactor for eNOS activation ^47^; decreasing L-arginine bioavailability^48^ limiting the substrate for NO production; impairing phosphorylation patterns promoting eNOS deactivation ^49^; and inducing oxidative inactivation of NO. ^50^ The near-restoration of endothelium-dependent vasodilation during acute antioxidant administration supports future vitamin C supplementation intervention studies as a viable intervention strategy aimed at improving vascular health after SCI.

There are a few experimental considerations to note. Firstly, Given the cross-sectional design of the present study, we cannot discount the possibility that genetic and/or lifestyle behaviors may have influenced our results. However, to minimize lifestyle behaviors known to influence endothelial function, we studied sedentary adults who were nonsmokers and were not taking any medication or supplements that could influence endothelial vasomotor function. In addition, to isolate the primary influence of SCI, all subjects were free of other cardiometabolic abnormalities such as hypertension, dyslipidemia, obesity, and type 2 diabetes. Secondly, we did not perform the necessary eNOS inhibition studies to confirm reduced eNOS activity as a central mechanism underlying the observed SCI-related impairment in vasodilation in response to both acetylcholine and isoproterenol. In addition, our study was not designed to isolate the source(s) of oxidative stress related to SCI. Finally, SCI populations are heterogeneous and the current study focuses primary on paraplegia. Differences in level of injury and severity of injury may influence both vascular endothelial function and the role of oxidative stress. Future studies are needed to address this important issue.

From a clinical perspective, it is important to emphasize that our study cohort of adults with SCI were free of overt cardiovascular disease and traditional cardiometabolic risk factors, allowing us to clinically isolate the effects of SCI *per se* on endothelial function. Our finding of SCI-related endothelial vasodilator dysfunction supports, and provides potential mechanistic insight for, population studies demonstrating that individuals with SCI experience disproportionately higher rates of major adverse cardiovascular events and greater atherosclerotic burden, even after adjustment for traditional risk factors. This suggests the presence of SCI-specific mechanisms that are not fully captured by conventional risk assessment, and highlights a critical gap that standard screening approaches derived from the general population may underestimate cardiovascular risk in SCI. This limitation is particularly relevant given that most individuals with SCI also accumulate traditional risk factors over time, compounding overall ASCVD risk.^51^

In conclusion, the results of the present study demonstrate that SCI resulting in paraplegia is associated with profound impairment in endothelium-dependent vasodilation. Endothelial vasodilator dysfunction likely contributes to the accelerated rate of atherosclerosis and atherosclerotic vascular events reported in individuals living with SCI. In addition, oxidative stress appears to be a central factor underlying SCI-related endothelial vasodilator dysfunction. Future studies are needed to determine the efficacy of therapeutic interventions, either lifestyle or pharmacologic, for improving endothelial function to mitigate the elevated ASCVD risk after SCI.

## ACKNOWLEDGEMENTS

We would like to thank all the subjects who participated in the study.

## SOURCES OF FUNDING

This study was supported, at least in part, by the American Heart Association award 24CDA1268, American Heart Association award 24TPA1301309, Craig H. Neilsen Foundation Spinal Cord Injury Research on the Translational Spectrum Award 1341523, Paralyzed Veterans of American Research Foundation PVA-3192 and PVA-3205, Craig Hospital Foundation 2821.

## DISCLOSURES

Andrew J. Park, Christopher A. DeSouza, Genevieve Madera, Clare Morey, Megan Summers, Vinicius P. Garcia, Auburn R. Berry, Samuel Ruzenne, Noah DeSouza, Joshua Holzer, Alexa Deitemeyer, Jared Greiner, and Brian L. Stauffer declare that they have no conflict of interest.

## Notes

### Competing Interest Statement

The authors have declared no competing interest.

## REFERENCES

1. Cragg JJ, Noonan VK, Krassioukov A, Borisoff J. Cardiovascular disease and spinal cord injury: results from a national population health survey. Neurology. Aug 20 2013;81(8):723–8. doi:10.1212/WNL.0b013e3182a1aa68

2. Myers J, Lee M, Kiratli J. Cardiovascular disease in spinal cord injury: an overview of prevalence, risk, evaluation, and management. Am J Phys Med Rehabil. Feb 2007;86(2):142–52. doi:10.1097/PHM.0b013e31802f0247

3. Yoo JE, Kim M, Kim B, et al. Increased Risk of Myocardial Infarction, Heart Failure, and Atrial Fibrillation After Spinal Cord Injury. J Am Coll Cardiol. Feb 20 2024;83(7):741–751. doi:10.1016/j.jacc.2023.12.010

4. Lee CS, Lu YH, Lee ST, Lin CC, Ding HJ. Evaluating the prevalence of silent coronary artery disease in asymptomatic patients with spinal cord injury. Int Heart J. May 2006;47(3):325–30. doi:10.1536/ihj.47.325

5. Matos-Souza JR, Pithon KR, Ozahata TM, Gemignani T, Cliquet A, Jr., Nadruz W, Jr. Carotid intima-media thickness is increased in patients with spinal cord injury independent of traditional cardiovascular risk factors. Atherosclerosis. Jan 2009;202(1):29–31. doi:10.1016/j.atherosclerosis.2008.04.013

6. Matos-Souza JR, Pithon KR, Ozahata TM, et al. Subclinical atherosclerosis is related to injury level but not to inflammatory parameters in spinal cord injury subjects. Spinal Cord. Oct 2010;48(10):740–4. doi:10.1038/sc.2010.12

7. Orakzai SH, Orakzai RH, Ahmadi N, et al. Measurement of coronary artery calcification by electron beam computerized tomography in persons with chronic spinal cord injury: evidence for increased atherosclerotic burden. Spinal Cord. Dec 2007;45(12):775–9. doi:10.1038/sj.sc.3102045

8. Lieberman JA, Hammond FM, Barringer TA, et al. Comparison of coronary artery calcification scores and National Cholesterol Education program guidelines for coronary heart disease risk assessment and treatment paradigms in individuals with chronic traumatic spinal cord injury. J Spinal Cord Med. 2011;34(2):233–40. doi:10.1179/107902610X12923394765733

9. Davignon J, Ganz P. Role of endothelial dysfunction in atherosclerosis. Circulation. Jun 15 2004;109(23 Suppl 1):III27–32. doi:10.1161/01.CIR.0000131515.03336.f8

10. Ludmer PL, Selwyn AP, Shook TL, et al. Paradoxical vasoconstriction induced by acetylcholine in atherosclerotic coronary arteries. N Engl J Med. Oct 23 1986;315(17):1046–51. doi:10.1056/NEJM198610233151702

11. Ross R, Glomset JA. The pathogenesis of atherosclerosis (first of two parts). N Engl J Med. Aug 12 1976;295(7):369–77. doi:10.1056/NEJM197608122950707

12. Xu S, Ilyas I, Little PJ, et al. Endothelial Dysfunction in Atherosclerotic Cardiovascular Diseases and Beyond: From Mechanism to Pharmacotherapies. Pharmacol Rev. Jul 2021;73(3):924–967. doi:10.1124/pharmrev.120.000096

13. Luscher TF, Tanner FC, Tschudi MR, Noll G. Endothelial dysfunction in coronary artery disease. Annu Rev Med. 1993;44:395–418. doi:10.1146/annurev.me.44.020193.002143

14. Yasue H, Matsuyama K, Matsuyama K, Okumura K, Morikami Y, Ogawa H. Responses of angiographically normal human coronary arteries to intracoronary injection of acetylcholine by age and segment. Possible role of early coronary atherosclerosis. Circulation. Feb 1990;81(2):482–90. doi:10.1161/01.cir.81.2.482

15. Celermajer DS, Sorensen KE, Gooch VM, et al. Non-invasive detection of endothelial dysfunction in children and adults at risk of atherosclerosis. Lancet. Nov 7 1992;340(8828):1111–5. doi:10.1016/0140-6736(92)93147-f

16. Schachinger V, Britten MB, Zeiher AM. Prognostic impact of coronary vasodilator dysfunction on adverse long-term outcome of coronary heart disease. Circulation. Apr 25 2000;101(16):1899–906. doi:10.1161/01.cir.101.16.1899

17. Meidell R. Southwestern internal medical conference: endothelial dysfunction and vascular disease. American Journal of the Medical Sciences. 1994;307:378–389.

18. Luscher T, Tanner F, Tschudi M, Noll G. Endothelial dysfunction in coronary artery disease. Annual Reviews of Medicine. 1993;44:395–418.

19. Vita J, Keaney J, Loscalzo J. Endothelial dysfunction in vascular disease. In: Loscalzo J, Creager M, Dzau V, eds. Vascular Medicine A Textbook of Vascular Biology and Diseases. 2 ed. Little Brown and Company; 1996:245–265.

20. Mano T, Masuyama T, Yamamoto K, et al. Endothelial dysfunction in the early stage of atherosclerosis precedes appearance of intimal lesions assessable with intravascular ultrasound. Am Heart J. Feb 1996;131(2):231–8. doi:10.1016/s0002-8703(96)90346-4

21. Bauersachs J, Bouloumie A, Fraccarollo D, Hu K, Busse R, Ertl G. Endothelial dysfunction in chronic myocardial infarction despite increased vascular endothelial nitric oxide synthase and soluble guanylate cyclase expression: role of enhanced vascular superoxide production. Circulation. Jul 20 1999;100(3):292–8. doi:10.1161/01.cir.100.3.292

22. Cosentino F, Rubattu S, Savoia C, Venturelli V, Pagannonne E, Volpe M. Endothelial dysfunction and stroke. J Cardiovasc Pharmacol. Nov 2001;38 Suppl 2:S75–8. doi:10.1097/00005344-200111002-00018

23. Zuchi C, Tritto I, Carluccio E, Mattei C, Cattadori G, Ambrosio G. Role of endothelial dysfunction in heart failure. Heart failure reviews. Jan 2020;25(1):21–30. doi:10.1007/s10741-019-09881-3

24. de Groot PC, Poelkens F, Kooijman M, Hopman MT. Preserved flow-mediated dilation in the inactive legs of spinal cord-injured individuals. Am J Physiol Heart Circ Physiol. Jul 2004;287(1):H374–80. doi:10.1152/ajpheart.00958.2003

25. Kooijman M, Thijssen DH, de Groot PC, et al. Flow-mediated dilatation in the superficial femoral artery is nitric oxide mediated in humans. J Physiol. Feb 15 2008;586(4):1137–45. doi:10.1113/jphysiol.2007.145722

26. Thijssen DH, Kooijman M, de Groot PC, et al. Endothelium-dependent and -independent vasodilation of the superficial femoral artery in spinal cord-injured subjects. J Appl Physiol (1985). May 2008;104(5):1387–93. doi:10.1152/japplphysiol.01039.2007

27. Phillips AA, Krassioukov AV. Contemporary Cardiovascular Concerns after Spinal Cord Injury: Mechanisms, Maladaptations, and Management. J Neurotrauma. Dec 15 2015;32(24):1927–42. doi:10.1089/neu.2015.3903

28. Groah SL, Weitzenkamp D, Sett P, Soni B, Savic G. The relationship between neurological level of injury and symptomatic cardiovascular disease risk in the aging spinal injured. Spinal Cord. Jun 2001;39(6):310–7. doi:10.1038/sj.sc.3101162

29. Krum H, Howes LG, Brown DJ, et al. Risk factors for cardiovascular disease in chronic spinal cord injury patients. Paraplegia. Jun 1992;30(6):381–8. doi:10.1038/sc.1992.87

30. Rupp R, Biering-Sorensen F, Burns SP, et al. International Standards for Neurological Classification of Spinal Cord Injury: Revised 2019. Top Spinal Cord Inj Rehabil. Spring 2021;27(2):1–22. doi:10.46292/sci2702-1

31. DeSouza CA, Shapiro LF, Clevenger CM, et al. Regular aerobic exercise prevents and restores age-related declines in endothelium-dependent vasodilation in healthy men. Circulation. Sep 19 2000;102(12):1351–7. doi:10.1161/01.cir.102.12.1351

32. Boland R, Adams R. Development and evaluation of a precision forearm and hand volumeter and measuring cylinder. J Hand Ther. Oct-Dec 1996;9(4):349–58. doi:10.1016/s0894-1130(96)80041-x

33. Frei B, England L, Ames B. Ascorbate is an outstanding antioxidant in human blood plasma. Proceedings of the National Academy of Sciences, USA. 1989;86:6377–6381.

34. Taddei S, Virdis A, Ghiadoni L, et al. Age-related reduction of NO availability and oxidative stress in humans. Hypertension. 2001;38:274–279.

35. Shimokawa H. Primary endothelial dysfunction: atherosclerosis. J Mol Cell Cardiol. Jan 1999;31(1):23–37. doi:10.1006/jmcc.1998.0841

36. Iranami H, Hatano Y, Tsukiyama Y, Maeda H, Mizumoto K. A beta-adrenoceptor agonist evokes a nitric oxide-cGMP relaxation mechanism modulated by adenylyl cyclase in rat aorta. Halothane does not inhibit this mechanism. Anesthesiology. Nov 1996;85(5):1129–38. doi:10.1097/00000542-199611000-00022

37. Gray DW, Marshall I. Novel signal transduction pathway mediating endothelium-dependent beta-adrenoceptor vasorelaxation in rat thoracic aorta. Br J Pharmacol. Nov 1992;107(3):684–90. doi:10.1111/j.1476-5381.1992.tb14507.x

38. Taddei S, Virdis A, Ghiadoni L, Magagna A, Salvetti A. Vitamin C improves endothelium-dependent vasodilation by restoring nitric oxide activity in essential hypertension. Circulation. Jun 9 1998;97(22):2222–9. doi:10.1161/01.cir.97.22.2222

39. Van Guilder GP, Hoetzer GL, Greiner JJ, Stauffer BL, DeSouza CA. Acute and chronic effects of vitamin C on endothelial fibrinolytic function in overweight and obese adult humans. J Physiol. Jul 15 2008;586(14):3525–35. doi:10.1113/jphysiol.2008.151555

40. Lubos E, Handy DE, Loscalzo J. Role of oxidative stress and nitric oxide in atherothrombosis. Front Biosci. May 1 2008;13:5323–44. doi:10.2741/3084

41. Bastani NE, Kostovski E, Sakhi AK, et al. Reduced antioxidant defense and increased oxidative stress in spinal cord injured patients. Arch Phys Med Rehabil. Dec 2012;93(12):2223–8 e2. doi:10.1016/j.apmr.2012.06.021

42. Liu D, Liu J, Wen J. Elevation of hydrogen peroxide after spinal cord injury detected by using the Fenton reaction. Free Radic Biol Med. Aug 1999;27(3-4):478–82. doi:10.1016/s0891-5849(99)00073-8

43. Visavadiya NP, Patel SP, VanRooyen JL, Sullivan PG, Rabchevsky AG. Cellular and subcellular oxidative stress parameters following severe spinal cord injury. Redox Biol. Aug 2016;8:59–67. doi:10.1016/j.redox.2015.12.011

44. Zhang C, Zhai T, Zhu J, et al. Research Progress of Antioxidants in Oxidative Stress Therapy after Spinal Cord Injury. Neurochem Res. Dec 2023;48(12):3473–3484. doi:10.1007/s11064-023-03993-x

45. Yu M, Wang Z, Wang D, Aierxi M, Ma Z, Wang Y. Oxidative stress following spinal cord injury: From molecular mechanisms to therapeutic targets. J Neurosci Res. Oct 2023;101(10):1538–1554. doi:10.1002/jnr.25221

46. McMillan DW, Bigford GE, Farkas GJ. The Physiology of Neurogenic Obesity: Lessons from Spinal Cord Injury Research. Obes Facts. 2023;16(4):313–325. doi:10.1159/000530888

47. Fei J, Demillard LJ, Ren J. Reactive oxygen species in cardiovascular diseases: an update. Exploration of Medicine. 2022;3(2):188–204. doi:10.37349/emed.2022.00085

48. Janaszak-Jasiecka A, Ploska A, Wieronska JM, Dobrucki LW, Kalinowski L. Endothelial dysfunction due to eNOS uncoupling: molecular mechanisms as potential therapeutic targets. Cell Mol Biol Lett. Mar 9 2023;28(1):21. doi:10.1186/s11658-023-00423-2

49. Karaa A, Kamoun WS, Clemens MG. Oxidative stress disrupts nitric oxide synthase activation in liver endothelial cells. Free Radical Biology and Medicine. 2005/11/15/ 2005;39(10):1320–1331. doi:10.1016/j.freeradbiomed.2005.06.014

50. Förstermann U, Xia N, Li H. Roles of Vascular Oxidative Stress and Nitric Oxide in the Pathogenesis of Atherosclerosis. Circulation Research. 2017/02/17 2017;120(4):713–735. doi:10.1161/CIRCRESAHA.116.309326

51. Gater DR, Jr., Farkas GJ, Tiozzo E. Pathophysiology of Neurogenic Obesity After Spinal Cord Injury. Top Spinal Cord Inj Rehabil. 2021;27(1):1–10. doi:10.46292/sci20-00067

